# Early exposure to Western Diet exacerbates visual outcomes in female mice

**DOI:** 10.1101/2024.11.27.625688

**Authors:** David Meseguer, Jessica Furtado, Thomas Zapadka, Amanda Rodriguez, Daxiang Na, Jose G. Grajales-Reyes, Jonathan Demb, Anne Eichmann, Marc Schneeberger

## Abstract

Obesity, a growing pandemic in Western societies, significantly impacts metabolic health and contributes to visual disorders. While the systemic consequences of obesity, such as chronic inflammation and insulin resistance, are well-studied in adults, its early-life effects on retinal health remain underexplored. Using a maternal Western Diet (WD) exposure model, we investigated the developmental impact of early-life metabolic disturbances on retinal and cognitive function. Our findings reveal that WD exposure from gestation to early adulthood accelerates the onset of features resembling diabetic retinopathy, including increased retinal vascularization, inflammation, and compromised blood-retina barrier integrity, observed within just four months. Females exhibited heightened vulnerability, showing pronounced ocular defects such as anophthalmia, microphthalmia, and congenital cataracts. These results underscore a critical developmental window during which metabolic disruptions predispose to sex-specific retinal and neurovascular pathologies. This work bridges the link between pediatric and adult obesity, highlighting the urgent need for early interventions to mitigate long-term visual impairments that could further impair recognition memory.

## Introduction

The etiology of obesity, the pandemic of modern societies, is multifactorial, involving genetic, environmental, and lifestyle contributors (**Bluher, 2019**). The systemic consequences of obesity, including chronic inflammation, insulin resistance, and cardiovascular complications, are well-documented (**Saltiel & Olefsky, 2017**). These features are the underlying factors of diabetic retinopathy, a metabolic complication that remains understudied in the context of pediatric obesity (**Antonetti et al., 2021**). Maternal obesity, diabetes, and hyperglycemia have been shown to predispose offspring to metabolic disorders, including retinal abnormalities, irrespective of genetic predisposition or maternal BMI (**Catalano & Shankar, 2017**).

Childhood obesity has become a critical global health issue, with recent data from the World Health Organization indicating that over 340 million children and adolescents aged 5–19 years and 39 million children under five were affected by overweight or obesity in 2022 (**WHO, 2023**). In the United States, the prevalence of childhood obesity has continued to rise, with the CDC reporting a 21.1% obesity rate and a 7.0% severe obesity rate among individuals aged 2–19 years in 2023 (**CDC, 2023**). This trend has been exacerbated by factors such as the COVID-19 pandemic, particularly among younger children, emphasizing the urgent need for effective interventions (**Rundle et al., 2020**).

Emerging research reveals links between childhood obesity and microvascular alterations in several ocular layers, including the retina (**Dezor-Garus J., 2023**). Dyslipidemia, insulin resistance, and non-alcoholic fatty liver disease (NAFLD) are known to disrupt retinal microvasculature. For instance, studies indicate a positive correlation between hepatic fibrosis in pediatric NAFLD and retinopathy signs, as well as elevated triglycerides, basal insulin, and HOMA-IR levels in children with retinopathy compared to those without (**Pacifico et al., 2020**). Additionally, high insulin-like growth factor 1 (IGF-1) levels have been implicated in retinal microvascular damage, particularly in overweight and obese children (**Travers et al. 1998**). These findings underscore the role of hyperinsulinemia, inflammation, and hormonal dysregulation in microvascular pathogenesis.

Microvascular changes in obese children, such as retinal venular dilatation and arterial narrowing, may reflect systemic endothelial dysfunction and inflammatory processes (**Dezor-Garus J., 2023**). Factors like leptin, which impairs endothelium-dependent vasodilation, and increased blood volume in obesity further contribute to these vascular alterations (**Stanek A, 2021**). However, the precise mechanisms connecting obesity to retinal vessel morphology remain unclear and warrant further exploration.

In animal models, prolonged WD exposure has been shown to induce retinal defects after nine to twelve months (**Keeling E et al. 2022**). However, differences in retinal vascular development between humans and mice necessitate alternative approaches to studying early-life influences when the vascular endothelium of central circuits is more vulnerable. The retinal vasculature in mice develops postnatally, making maternal WD exposure an intriguing model for understanding the impact of early-life metabolic disturbances on retinal health (**Selvam et al. 2018**).

Our research demonstrates that maternal WD exposure induces key features of diabetic retinopathy—such as inflammation, increased vascularization, and blood-retina barrier (BRB) leakage—within just four months of dietary exposure, only in female mice. Moreover, in females, we also observed macroscopic ocular defects (anophthalmia, microphthalmia, and cataracts), establishing a novel sexual dimorphic model to investigate the link between obesity and retinal abnormalities. This work offers a new lens to visualize the molecular underpinnings of diabetic retinopathy and highlights the potential of targeting early-life metabolic disturbances to mitigate obesity-associated retinal pathologies.

## Results

### Western Diet exposure during early life results into a sexual dimorphic energy homeostasis phenotype

To better represent a human environment of childhood obesity and evaluate the impact of Western Diet exposure from early development into adolescence in mice, we provided female C57Bl/6 virgin mice with a WD (high-fat/high-sucrose) starting from gestation through four months of age (**Figure 1A**). Of note, at the day of birth (DOB), litter size was adapted to 7-8 pups per mother, ensuring a similar nutritional environment for each litter and avoiding metabolic disorders associated with a small litter size. The offspring was divided in different cohorts to provide the same age between animals, and we monitored the weight during the exposure to the WD. Interestingly, while females fed with a WD showed resistance, maintaining comparable weights to those on a standard chow diet, consistent with prior observations **(Vogt et al., 2014)**, males exhibited a distinct overweight phenotype (**Figure 1B**). While both genders exhibited higher food intake and reduced core body temperature (**Figure 1C-D**) sex-specific difference extended to other metabolic parameters, with males showing increased adiposity than females (**Figure 1E**). Despite these differences in weight and adiposity, both males and females demonstrated hyperglycemia, consistent with the prediabetic phenotype typically associated with early WD-fed animals **(Samuelsson et al., 2008)**. Moreover, gestationally exposed WD mice exhibited glucose intolerance and high glucose levels compared to littermate standard chow diet fed controls (**Figure 1F-G**).

**Figure 1:**
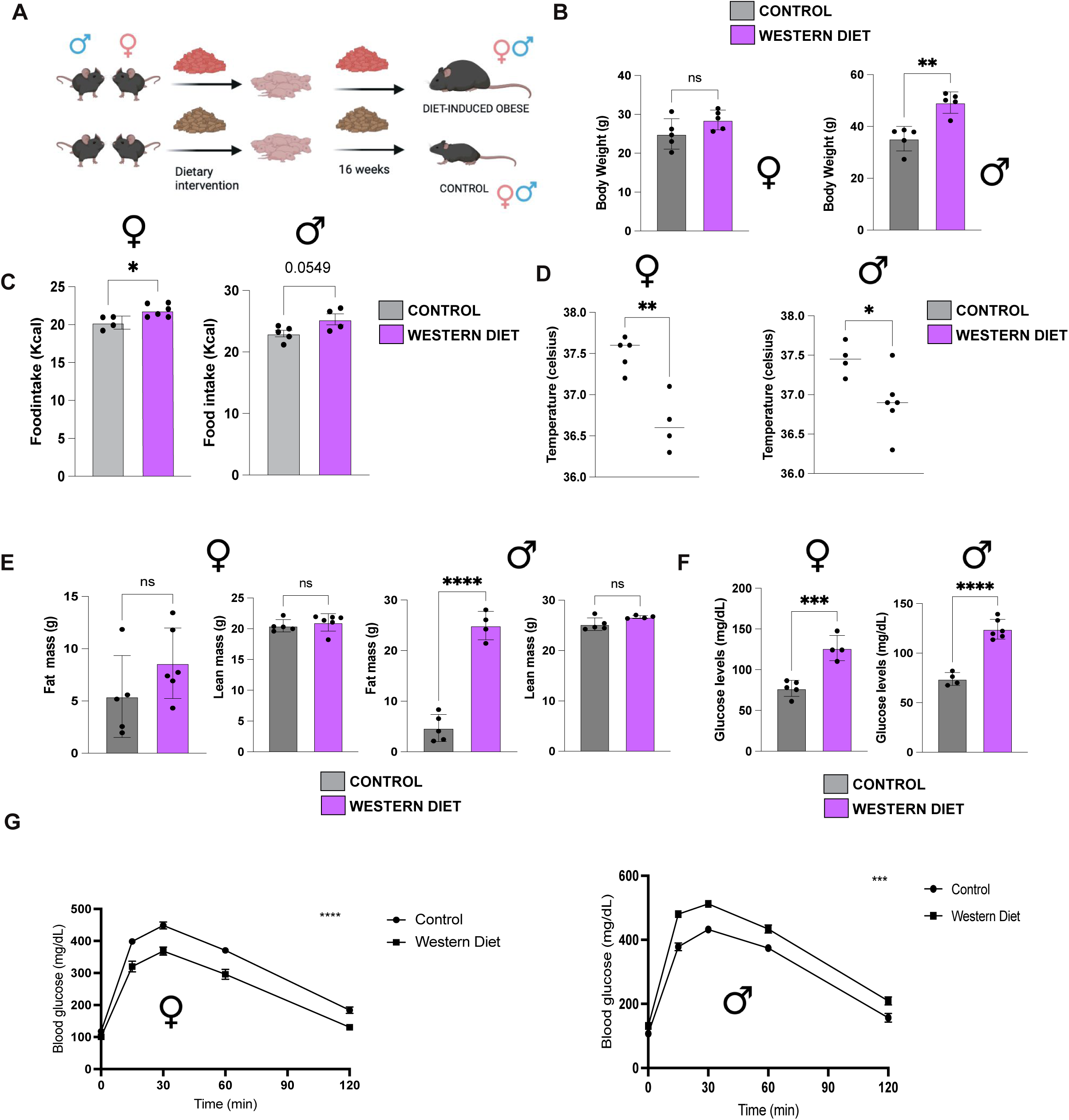
Early exposure to WD differentially affects physiological parameters of the offspring. **A)** Breeding scheme and diet intervention to impact all central nervous system by a dietary shift towards WD. **B)** Body weight (n= 4-6/group) in males and females after 16 weeks exposure to WD from embryonic to adult stages of life. **C)** Lean mass and fat mass in grams of standard diet and WD fed animals after 16 weeks exposure to from embryonic to adult stages of life (n= 4-6). **D)** Daily food intake and **E)** anal core body temperature recorded at 7a.m. in standard diet and WD fed animals after 16 weeks exposure to from embryonic to adult stages of life (n =4-6). **F)** Fasted blood glucose after a 16-hour fasting and **G)** glucose tolerance test after 4 hours fasting (glucose = 2g/kg) in standard diet and WD-WD fed animals after 16 weeks exposure to from embryonic to adult stages of life (n =4-6). Data are expressed as mean ± SEM. \**P*<0.05; \*\**P*<0.01; \*\*\**P*<0.001.; ****p<0.0001; ns: not statistically significant.

These findings underscore the complex, sex-specific metabolic effects of prolonged Western Diet exposure, emphasizing the importance of studying both sexes to capture the full spectrum of obesity-related pathophysiology.

### The environment associated with Western Diet exposure underlies an enhanced predisposition to macroscopic ocular defects

Anophthalmia, microphthalmia, and congenital cataracts reflect disruptions in early eye development, often linked to defects in optic vesicle formation, lens induction, or retinal differentiation. These anomalies, from absent or underdeveloped eyes to lens opacities at birth, highlight critical molecular pathways in ocular embryogenesis and are frequently associated with genetic syndromes or environmental exposures during gestation. Black six mice occasionally exhibit such deficiencies (https://www.jax.org/news-and-insights/1995/october/microphthalmia-and-ocular-infections-in-inbred-c57-black-mice)

To explore these effects, we investigated the offspring of maternal Western Diet-fed mice in 3 independent cohorts to rule out any genetic predisposition present in the breeders (**Figure 2A**). Strikingly, 40% of female offspring exhibited macroscopic ocular abnormalities, including anophthalmia (6%), microphthalmia (13%), and congenital cataracts (20%). These abnormalities exhibited a pronounced sex-specific pattern, with an incidence of 40 % in females versus 3 % in males (**Figure 2B**).

**Figure 2:**
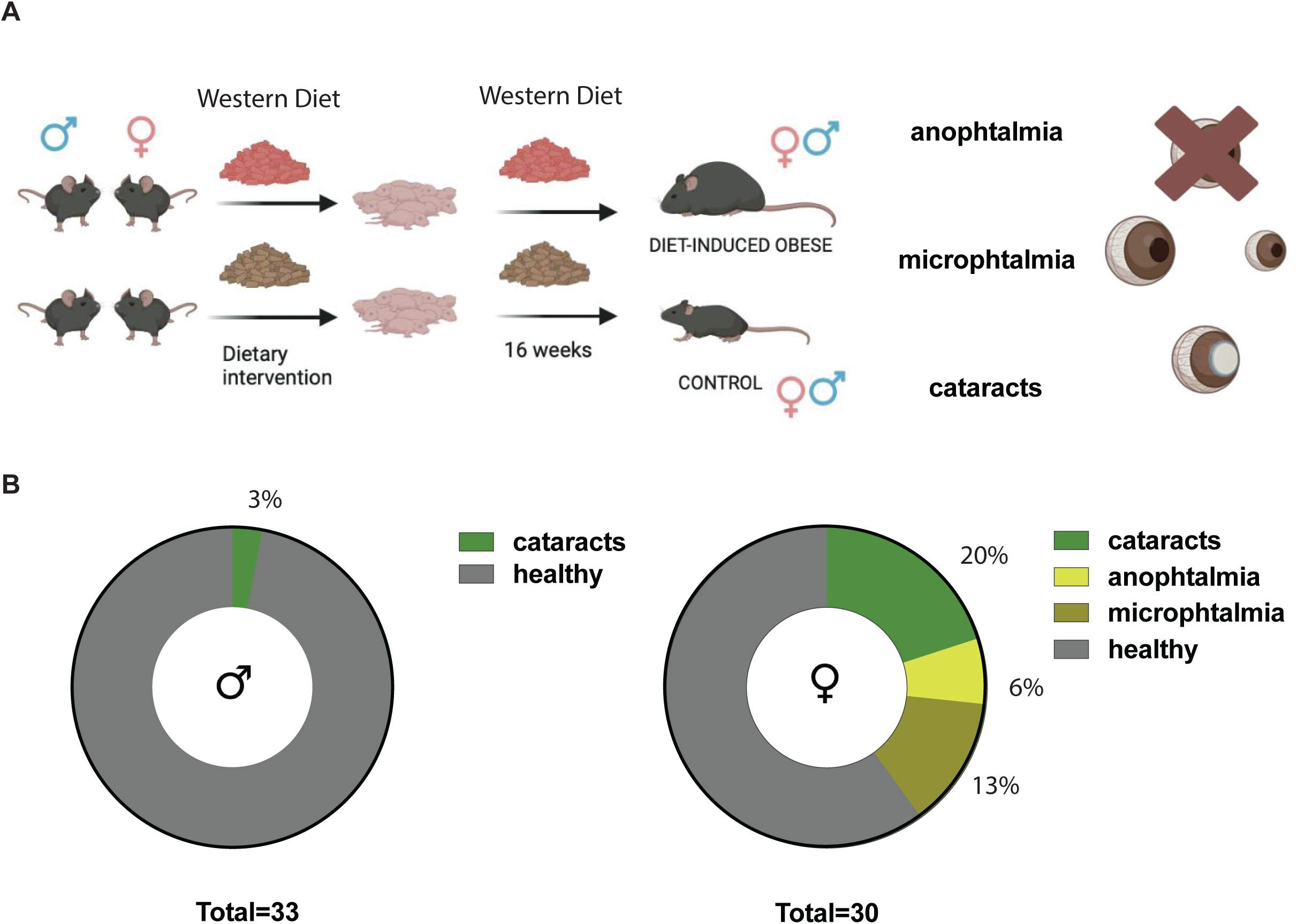
Early exposure to WD induces gender specific macroscopic changes in the visual system. **A)** Breeding scheme and associated eye disorders from being exposed to WD during developmental stages. **B)** Quantification of the percentage of male and female mice showcasing macroscopic vision problems due to exposure to WD during development. Strikingly females exhibit a more profound effect. Animals obtained from three different independent crosses of 6 female mice.

These findings suggest that early exposure to a WD provides a valuable model for studying the molecular pathways contributing to the higher prevalence of visual disorders observed in women than men **(AAO 2024)**. This model may help to identify sex-specific therapeutic targets and interventions for congenital and metabolic-related visual disorders.

### Western Diet exposure during early life results into retinovascular defects

The nervous system’s microvasculature undergoes critical development postnatally, rendering it highly sensitive to environmental influences during early life **(Rice et al. 2000)**. Exposure to a WD during this vulnerable period increases inflammatory cues, which can disrupt neurovascular endothelium, a key regulator of blood-retinal barrier integrity **(Rudraraju et al. 2020)**. Since retinal vascular integrity is crucial for maintaining normal retinal function **(Eltanani 2022)**, we examined structural and functional alterations in neurovascular physiology in WD-exposed mice compared to littermate controls during development.

To evaluate the impact of nutritional changes during development on retinal vasculature in mice without macroscopic alterations in the visual compartment, we dissected the retinas from these animals and performed immunofluorescence staining using Isolectin B4 (IB4), a marker of endothelial cells **(Boyé 2022)**. This approach allowed us to quantify the total retinal vasculature by measuring the area and mean of IB4-positive cells. Interestingly, we observed a noticeable trend toward increased vascular density in females exposed to a WD during childhood (**Figure 3A**), though this effect did not reach statistical significance (p = 0.06). These seem to be in the same direction as the human data, showing that neovascularization is one of the main pathophysiological signs of DR. In contrast, no such trend was evident in males, highlighting a potential sex-specific response to Western Diet exposure.

**Figure 3:**
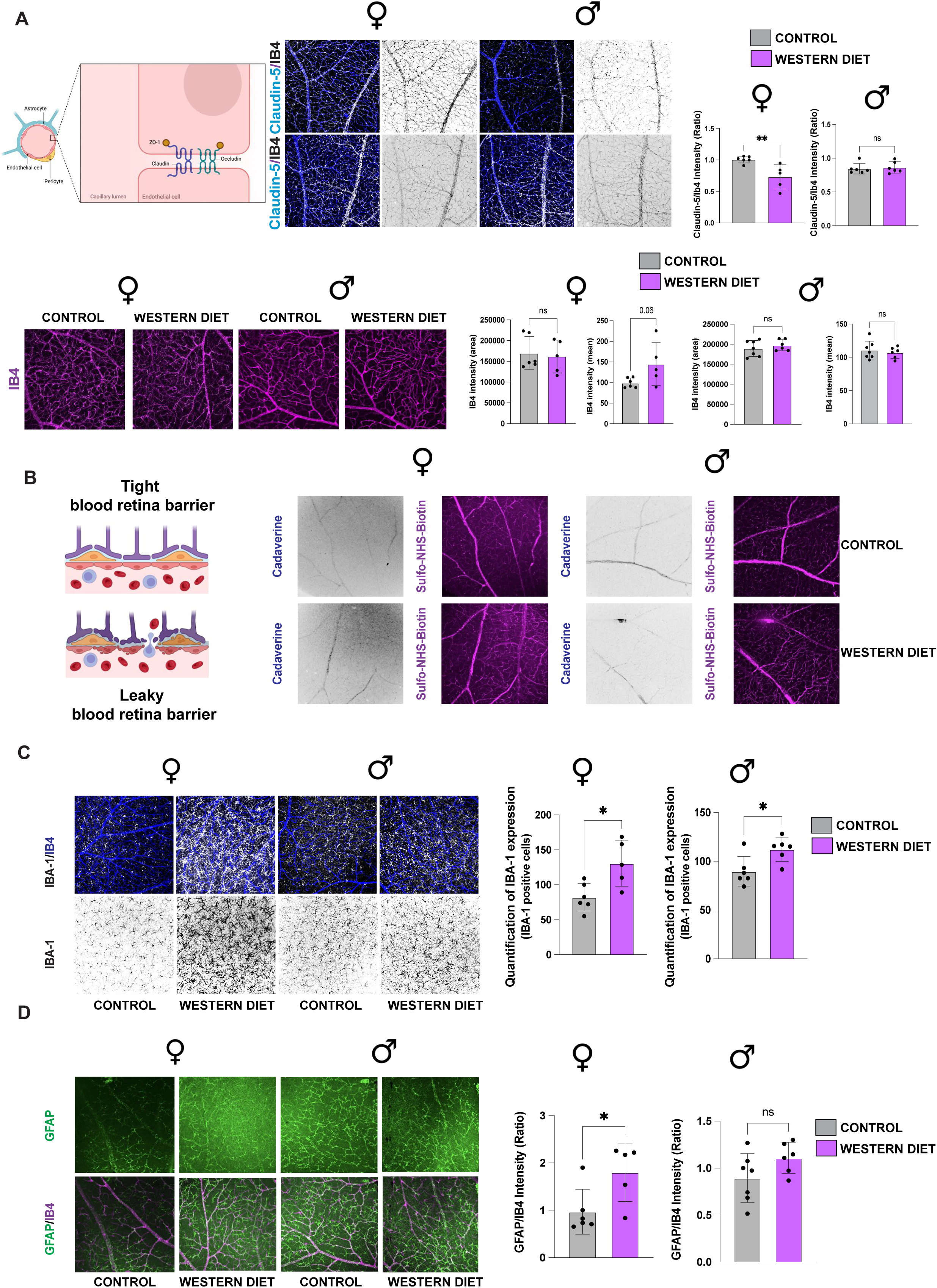
Early exposure to WD induces neurovascular and neuroinflammatory response in the mouse retina in a sexually dimorphic manner. **A)** Expression of claudin-5 normalized by vascular density (IB4) of individual petals of retinas from control and WD fed animals. **B)** Immunofluorescence staining and confocal imaging on individual petals, 30 minutes after retro-orbitally injection of 5mg of Cadaverine per gram of body weight and 1mg of sulfo-NHS-biotin/ mice in the case of the controls and 1.5 mg/ mice in the case of the obese animals. Qualitative results showing areas of leak. **C)** Immunofluorescence staining of IBA-1+ cells and confocal imaging of individual petals of retinas from control and WD fed animals **D)** Immunofluorescence staining of GFAP+ cells and confocal imaging of individual petals of retinas from control and WD fed animals. Quantification of superficial GFAP+ cells normalized by vascular density (IB4). All data are shown as mean+/-SEM. Two group-one factor comparisons were performed using a two-tailed unpaired Student’s t test. Symbols used are: *p < 0.005; **p < 0.001

Structural alterations in retinal vasculature are intimately related to modifications in BRB integrity **(Kim 2023, Yao 2024).** To further investigate the impact of WD exposure during childhood on the integrity of the BRB, we employed a dual-method approach. First, we assessed the tight junction protein claudin-5 expression, a critical component of endothelial cell barrier integrity **(Boyé 2022)**. Second, we examined the leakage of two retro-orbitally injected dye tracers of 1 kDa molecular size: sulfo-NHS-biotin and cadaverine **(Li 2022)**. Strikingly, a significant reduction in claudin-5 expression was observed in females exposed to the WD in early life (**Figure 3A**) but not in males, indicating compromised tight junction integrity in only one of the sexes. This reduction was accompanied by notable regions of leakage of sulfo-NHS-biotin and cadaverine tracers into the retinal parenchyma (**Figure 3B**), suggesting increased permeability of the BRB. These findings highlight sex-specific vulnerabilities in BRB integrity in response to early-life nutritional changes.

The observed compromise in BRB integrity, particularly in females exposed to a Western Diet, could be caused by an inflammatory response, a potential underlying mechanism previously shown in different studies. To confirm the existence of an inflammatory environment in these animals, we next assessed retinal microglia, a hallmark of neuroinflammation, using immunostaining for IBA-1, a specific marker for microglia **(Noailles 2014)**. Microglia, as resident immune cells of the retina, play a crucial role in maintaining tissue homeostasis apart from the development of the retina during childhood **(Noailles 2014)**. IBA-1 staining following early-life Western Diet exposure revealed a significant increase in activated microglia in both female and male mice (**Figure 3C**). Of note, microglial morphology in early WD-exposed mice suggested heightened activation, as evidenced by a transition from ramified structures, indicative of a resting state, to a more amoeboid shape, characteristic of an activated state. This morphological shift reflects increased inflammatory activity, aligning with the observed retinal inflammation and compromised neurovascular integrity in WD-exposed females. Microglia activation could further contribute to retinal dysfunction, implicating neuroinflammatory pathways in the observed retinal and vascular abnormalities.

Upon early-life WD exposure, microglia-astrocyte crosstalk may be particularly relevant. The observed increase in activated microglia could lead to heightened astrocyte reactivity, further compromising the BRB and contributing to the structural and functional abnormalities identified in the retina. In the retina, microglia and astrocytes interact within the neurovascular unit, contributing to regulating blood-retinal barrier integrity and neuronal health. Microglia release pro-inflammatory cytokines such as TNF-α and IL-1β upon activation, stimulating astrocytes to adopt a reactive state. To assess the state of astrocytes and Muller glial cells in mouse retina, we conducted an immunofluorescence test on the glial fibrillary acidic protein (GFAP) marker. Quantification of GFAP in mouse retinas shows an increased astrocytic coverage after exposure to WD early in life in female but not in male mice (**Figure 3D**). This interplay underscores the significance of maintaining retinal homeostasis and the potential for dietary-induced disruptions to trigger a cascade of pathological inflammation that compromises BRB integrity.

Taken together, these results highlight the profound impact of early-life Western Diet exposure on retinal neurovascular health, mainly through sex-specific mechanisms. The findings demonstrate that WD exposure during critical developmental windows induces significant structural and functional changes in the retinal vasculature, with females showing increased vascularization and compromised BRB integrity. Reduced claudin-5 expression and selective permeability to small molecular tracers underscore the vulnerability of endothelial tight junctions to dietary influences. The observed microglial activation in both sexes further implicates inflammatory pathways in the disruption of retinal homeostasis. These activated microglia may contribute to astrocytic reactivity, as increased GFAP expression indicates, suggesting a dynamic interplay between glial cells that exacerbates BRB dysfunction. This glial crosstalk underscores the broader neuroinflammatory cascade triggered by early-life nutritional changes and its potential to drive retinal pathologies.

### Western Diet Exposure Does Not Impair Electroretinogram (ERG) Function After 4 Months of Feeding

To evaluate the functional impact of early-life exposure to a WD on retinal physiology in mice without macroscopic defects (**Figure 2**), we assessed retinal activity using electroretinography (ERG). This technique measures the electrical responses of retinal cells to light stimuli and, more specifically, the function of rods **(Hanke-Gogokhia et al., 2024).** In a scotopic ERG, which measures retinal responses under low-light conditions, the recorded waves reflect distinct layers of retinal activity. We quantified two components of the ERG: the scotopic a-wave and the scotopic b-wave. The a-wave represents the initial hyperpolarization of rod photoreceptors in response to light stimuli **(Hanke-Gogokhia et al. 2024)**. This negative deflection is a direct measure of photoreceptor function and provides insights into the health and responsiveness of these cells, which are critical for vision in dim lighting. The b-wave emerges as a positive deflection, signifying the activity of the inner retinal layers **(Hanke-Gogokhia et al. 2024)**. Specifically, it is generated by the depolarization of ON-bipolar cells and the involvement of Müller glial cells. The b-wave thus reflects the processing of visual signals transmitted from the photoreceptors to the inner retina. Together, the a- and b-waves comprehensively assess rod-mediated retinal function. Notably, scotopic ERGs in male and female mice exposed early to a Western Diet and their littermates fed a control diet showed no differences (**Figure 4A and 5A**). Neither quantification of the scotopic a-wave (**Figure 4B and 5B**) nor the scotopic b-wave (**Figure 4C and 5C**) showed significant differences.

**Figure 4:**
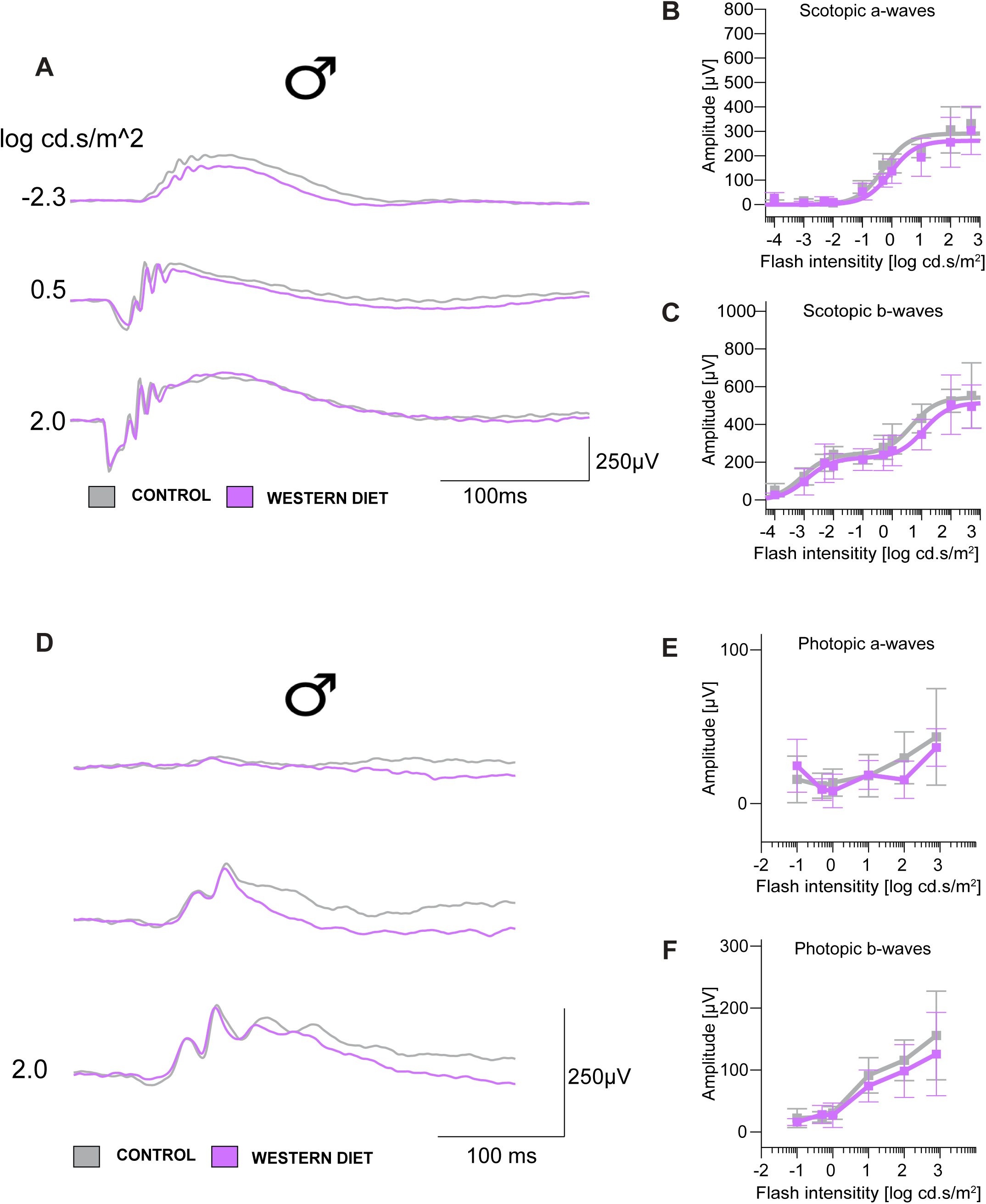
Scotopic and Photopic ERG responses are not altered in male mice fed to western diet. A) Average scotopic ERG responses (n = 5 animals per genotype) from control (black) and western diet (magenta) mice were recorded after 16 weeks of western diet. B) The a-wave C) and b-wave amplitudes are plotted as a function of increasing flash intensity (mean ± SD). Scotopic ERG responses from western diet animals were comparable to their littermate controls (black). D) Average photopic ERG responses. E) Amplitudes of photopic a-wave F) and b-wave were plotted over increasing intensity of single-flash stimuli. Cone-driven ERG b-waves recorded from both groups are normal. Comparisons reflect average response over two flash intensities within the gray bars; *p < 0.005; **p < 0.001; not statistically significant.

**Figure 5:**
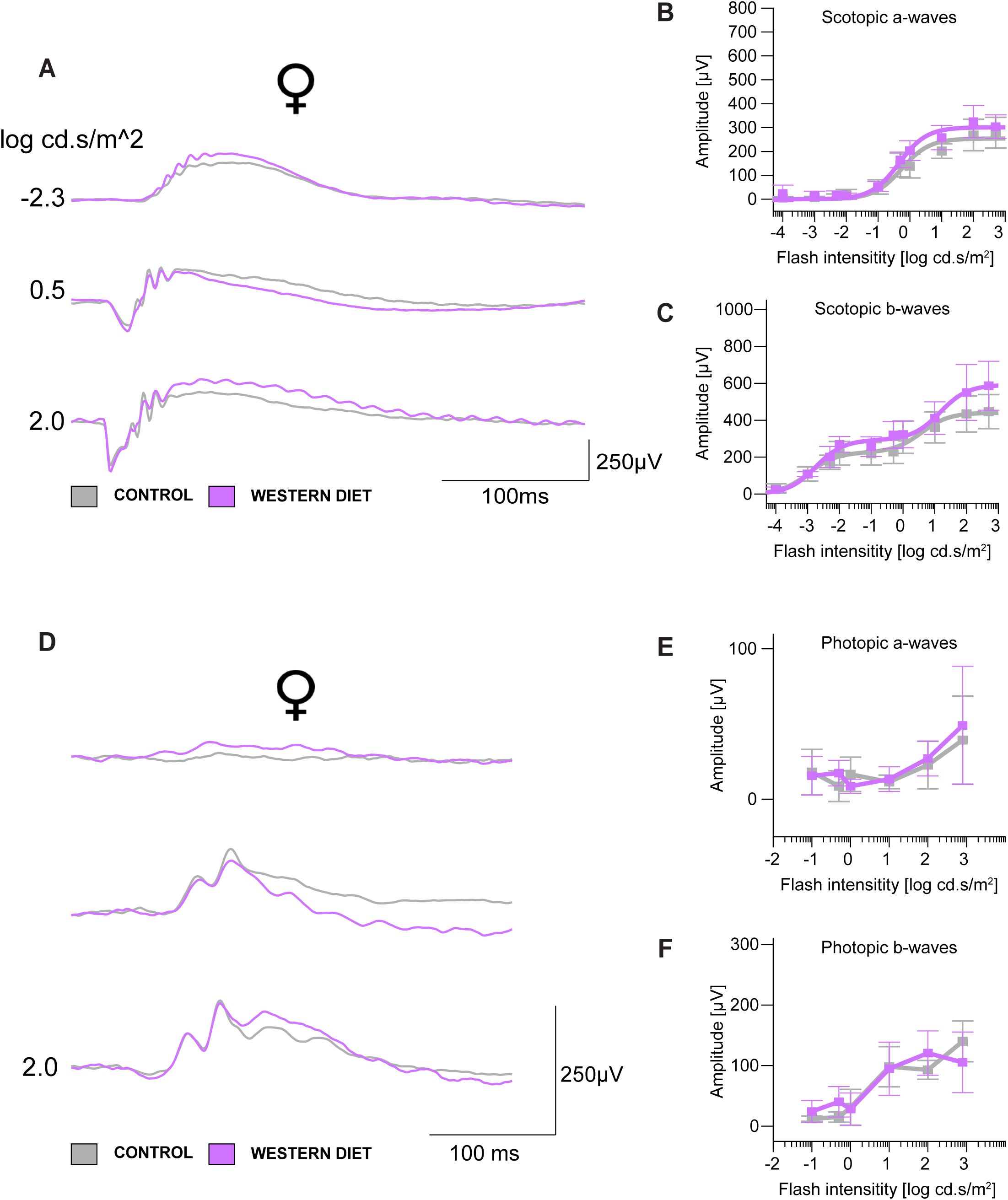
Scotopic and Photopic ERG responses are not altered in female mice fed to western diet. A) Average scotopic ERG responses (n = 5 animals per genotype) from control (black) and western diet (magenta) mice were recorded after 16 weeks of western diet. B) The a-wave C) and b-wave amplitudes are plotted as a function of increasing flash intensity (mean ± SD). Scotopic ERG responses from western diet animals were comparable to their littermate controls (black). D) Average photopic ERG responses. E) Amplitudes of photopic a-wave F) and b-wave were plotted over increasing intensity of single-flash stimuli. Cone-driven ERG b-waves recorded from both groups are normal. Comparisons reflect average response over two flash intensities within the gray bars; *p < 0.005; **p < 0.001; ns: not statistically significant.

Next, we performed a photopic ERG to assess the functional response of cone photoreceptors and their associated pathways in the retina under bright-light conditions. The photopic ERG isolates cone activity using intense background illumination to suppress rod responses. This approach focuses on the visual processes essential for color vision and visual acuity. In those conditions, both the a-wave, which represents the initial response of cone photoreceptors to light and marking the beginning of the visual signal, and the b-wave, originating from the ON-bipolar cells and Müller cells and reflecting the transmission and early processing of the visual signal within the retinal circuitry, showed comparable values between males and females after early exposure to WD or Standard Diet (**Figure 4 D-F and 5 D-F).**

Taken together, despite the structural and vascular alterations observed in retinal development, no significant impairment was detected in ERG recordings after 4 months of WD feeding. Both a-wave and b-wave amplitudes, which correspond to photoreceptor and bipolar cell responses, respectively, remained comparable between WD-fed mice and their littermate controls across all tested light intensities.

These findings suggest that while WD exposure induces microvascular and neuroinflammatory changes in the retina, the functional responses of the primary retinal circuitry to light stimulation are preserved at this stage.

### Impact of Early High-Fat Diet Exposure on Visually Related Cognitive Performance

Retinal dysfunction in offspring of mice exposed to a Western Diet during development, marked by compromised blood-retinal barrier integrity, heightened microglial activation, and increased astrocyte reactivity, does not significantly alter scotopic or photopic ERG parameters (**Figure 3-4**). However, these subtle changes may still impair overall visual perception. Impaired sensory input from retinal dysfunction, combined with broader neuroinflammatory processes and synaptic disturbances associated with WD exposure, could contribute to deficits in recognition memory and task performance.

Visual dysfunctions may extend to cognitive performance, particularly in tasks like the novel object recognition (NOR) test, which relies heavily on intact visual processing and memory to distinguish between familiar and novel objects **(Antunes et al. 2012).** Both genders showed a decrease in the total exploration time during the test. Interestingly, female mice exposed to a WD during early development displayed a diminished ability to recognize new objects, as they showed decrease frequency and cumulative duration with the novel object than standard-diet-fed controls. Although not statistically significant, we observed a trend in the discrimination index. (**Figure 6A**).

**Figure 6:**
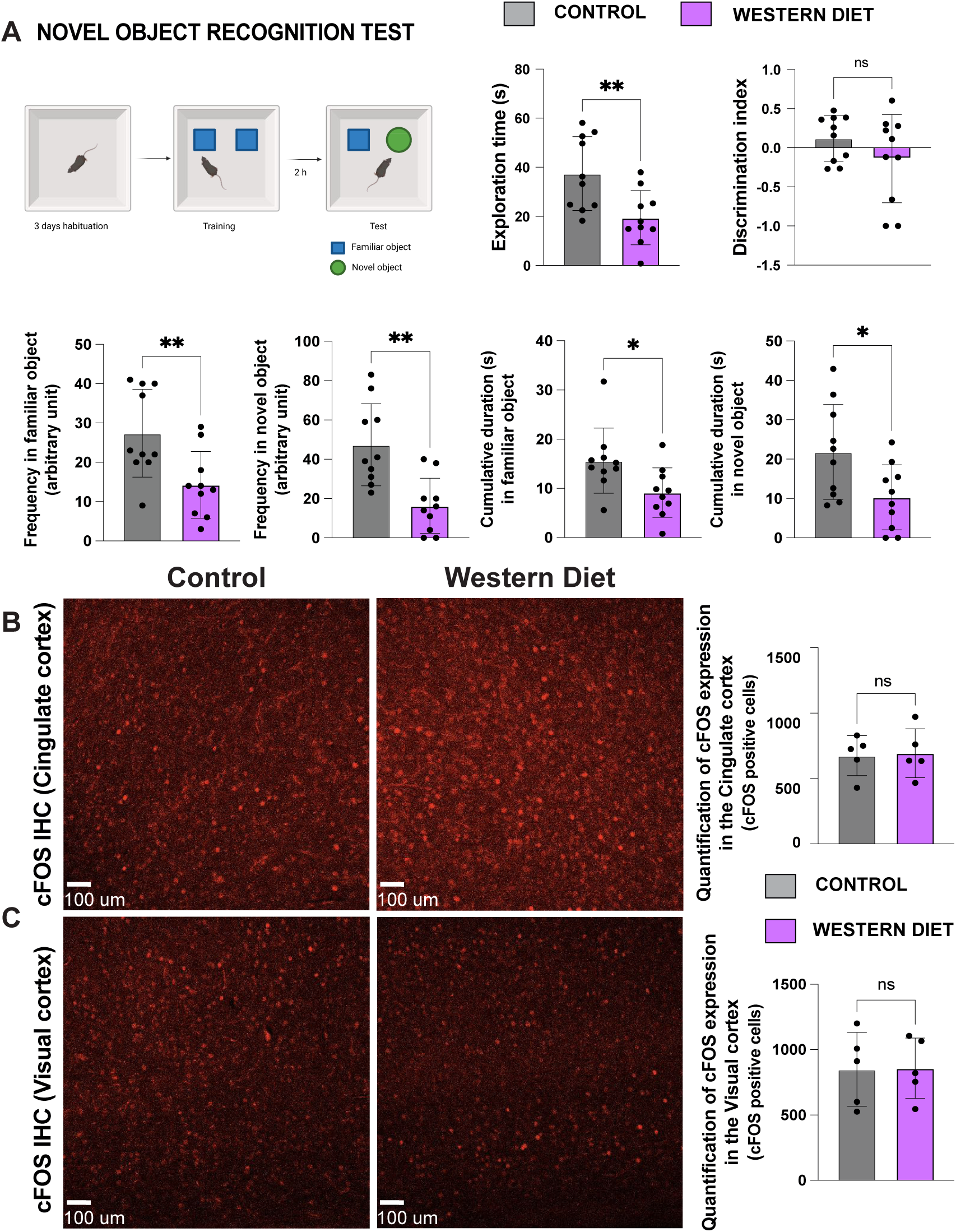
Exposition to western diet in early stages of development causes cognitive impairments but no changes in cFOS activation in somatosensory brain regions. **A)** Schematic illustration of the novel object recognition test (NORT). Recorded parameters to assess NORT performance in mice fed with either chow or WD during the test phase: exploration time, discrimination index (time exploring novel object + time exploring familiar object)/(time exploring novel object + time exploring familiar object), frequency in familiar object, frequency in novel object, cumulative duration in familiar object and cumulative duration in novel object. B) cFOS immunostaining in cingulate cortex (A24a) C) cFOS immunostaining in visual cortex (V1). cFOS staining was quantified as number of cFOS positive cells. Symbols used are: *p < 0.005; **p < 0.001; ns: statistically not significant.

To explore whether this impairment stemmed from deficits in memory circuits or visual processing regions, we assessed the expression of early activity markers, such as cFOS, in the medial prefrontal cortex and visual cortex. Surprisingly, no significant changes in cFOS expression were detected in these areas (**Figures 6B and 6C**), suggesting that alterations in these specific regions might not account for the observed NOR deficits. Future investigations combining advanced techniques, such as calcium imaging to track neuronal activity in real-time and transcriptomic analyses to evaluate gene expression changes, could elucidate whether the observed NOR deficits are primarily linked to disruptions in visual processing pathways or impairments in memory-related brain regions.

To further explore the potential impact on memory, we performed additional behavioral assays. The Barnes maze test, known for assessing spatial learning and memory, provides an opportunity to investigate spatial memory deficits in these animals directly **(Rodriguez Peris 2024)**. While female mice exposed early to the WD still recognized the target hole, they tended to explore less the rest of the maze, something we interpreted as a lack of motivation (**Figure 7A**). Additionally, we conducted the open field test (OFT) and elevated plus maze (EPM) to assess anxiety and exploratory behaviors-cognitive and emotional processes that are frequently impaired in obesity **(Keleher et al. 2018, Tsan et al. 2021)**. These tests provided further insights into the extent to which these behaviors are disrupted in female mice exposed early to a WD. Notably, female mice exposed to the WD during development exhibited reduced velocity and distance traveled and less time spent in the center of the OFT (**Figure 7B**). Similarly, in the EPM, they spent less time in both the open and closed arms (**Figure 7C**). These findings suggest that early WD exposure may impair exploratory and anxiety-related behaviors in females, indicating broader disruptions in cognitive and emotional functions, which could be linked to the neuroinflammatory and metabolic changes induced by the diet. Such impairments align with previous studies showing that WDs can disrupt neurodevelopmental processes and cognitive function, including anxiety regulation **(Hayes et al. 2024, Lopez-Taboada et al. 2020)**.

**Figure 7:**
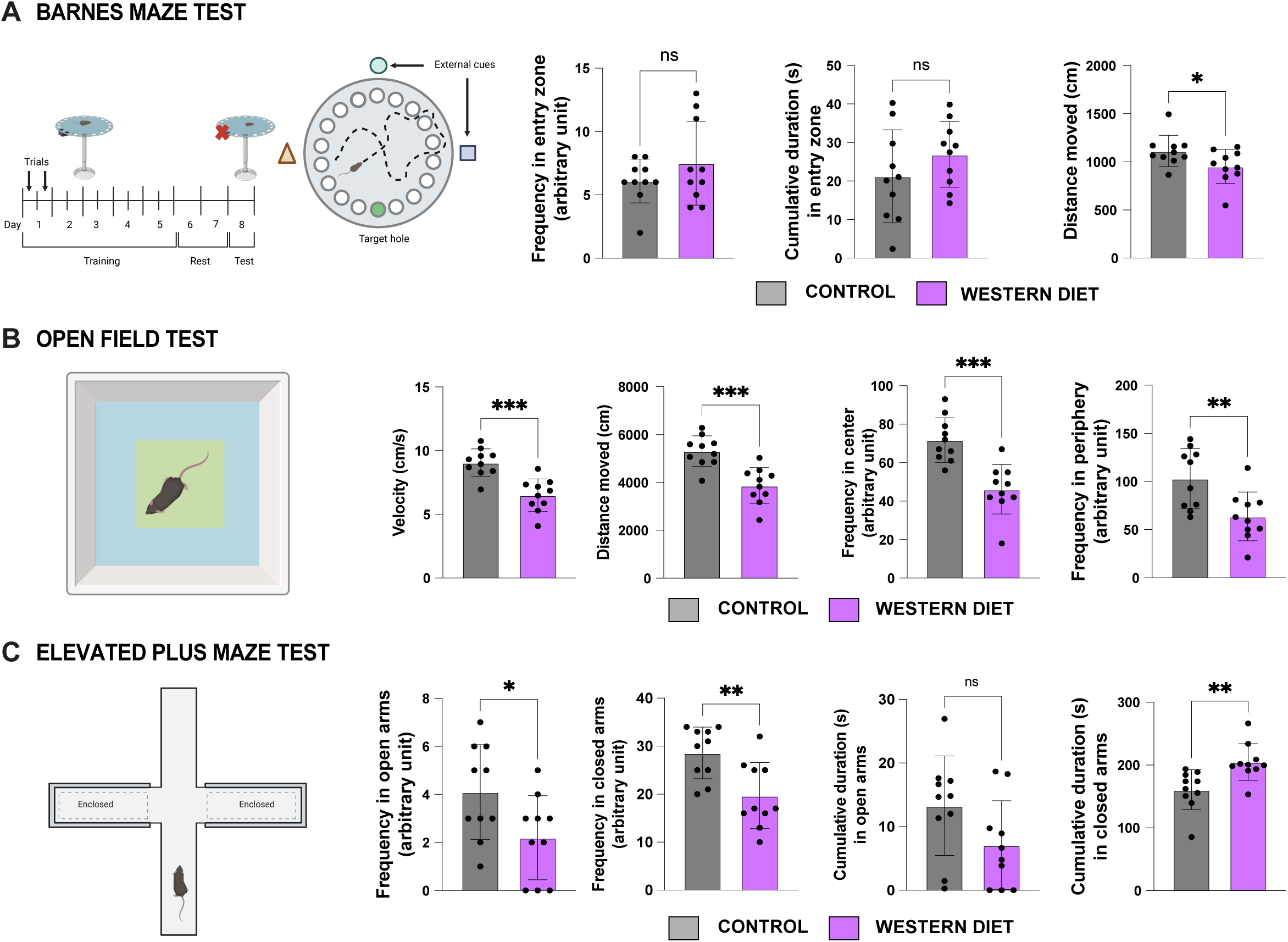
BMT, OFT and EP test show poor cognitive performance and increased levels of anxiety in animals fed with WD. **A)** Schematic illustration of BMT and experimental timeline. The green filled circle represents the scape hole and scape chamber location. The triangle, square and circle surrounded the maze represent the external clues. The position of the scape chamber remained constant on each trial. On the test day, the scape hole was closed and the chamber removed. Recorded parameters to assess BMT performance in mice fed with either chow or western diet for 4 months during the test phase: frequency in entry zone, cumulative duration in entry zone and total distance traveled. B) Schematic representation of OFT. Blue area represents the periphery while yellow area represents the center of the arena. Recorded parameters: velocity, distance moved, frequency in center and frequency in the periphery. C) Schematic representation of EPM. Analyzed parameters: Frequency in open arms, frequency in closed arms, cumulative duration in open arms and cumulative duration in closed arms. Symbols used are: *p < 0.005; **p < 0.001; ns: not statistically significant.

Our data demonstrate that early exposure to a WD induces a pro-inflammatory environment in the retina earlier than what it is typically seen in adults exposed to WDs after long-term exposure (12 months) **(Clarkson-Townsend 2021)**. This response presents with a notable sexual dimorphism, with females showing a more pronounced impact, mirroring trends observed in the human population, where women are more frequently affected by diet-related visual disorders such as diabetic retinopathy and age-related macular degeneration **(AAO 2024)**. The findings highlight the importance of early-life dietary influences in the development of retinal dysfunction and offer a promising model for investigating the pathophysiology of these diseases. Further mechanistic studies are necessary to explore the underlying molecular pathways driving these early-life diet-induced changes. This could lead to novel therapeutic strategies for visual impairments linked to metabolic dysfunction.

## Discussion

Childhood obesity poses a significant threat to the development of central neural circuits, with long-lasting consequences that extend into adulthood **(Logan et al. 2022)**. However, the full extent of these effects, particularly on processes like synaptic pruning, neurovascular development, and circuit plasticity, still needs to be understood. This gap in knowledge underscores the need for further investigation into how early metabolic challenges shape the brain’s structure and function over a lifetime. During critical periods of brain maturation, excessive adiposity and metabolic dysregulation can alter the delicate balance of neuroinflammatory processes, disrupt synaptic plasticity, and impair blood-brain barrier (BBB) integrity **(Feng et al. 2024)**. These changes can compromise the development of brain regions such as the hippocampus, prefrontal cortex, and hypothalamus, vital for cognition, emotional regulation, and metabolic homeostasis. For instance, several studies have shown that early-life obesity is associated with the existence of a neuroinflammatory environment highlighted by microglial activation and astrocyte dysfunction, leading to impaired learning, memory, and executive functioning **(Cope et al. 2018; Balasubramanian et al. 2020)**. Furthermore, systemic inflammation and insulin resistance resulting from obesity can exacerbate neural deficits, creating a vicious cycle that perpetuates cognitive and behavioral impairments throughout life **(Gómez-Apo et al. 2021)**. This inflammatory and metabolic burden also extends to other structures of the CNS such as the retina and optic pathways, disrupting neurovascular integrity and predisposing individuals to visual disorders such as diabetic retinopathy and age-related macular degeneration **(Kóvacs-Valasek 2023)**. These interconnected neural and ocular impairments highlight the urgent need for interventions targeting childhood obesity to preserve both brain and visual health across the lifespan. The present study investigates the impact of early-life WD exposure on retinal health, neurovascular integrity, and cognition, emphasizing the sexually dimorphic nature of these effects. Our findings build upon and extend existing literature regarding how early nutritional environments influence systemic and ocular health outcomes.

Our results demonstrate that female mice are more susceptible to WD-induced retinal and cognitive dysfunctions than males. This finding aligns with clinical studies documenting a higher prevalence of diet-related visual impairments in women, including diabetic retinopathy and age-related macular degeneration **(AAO 2024)**. This sex-specific vulnerability may stem from hormonal interactions with inflammatory and neurovascular pathways. For example, estrogen and other sex hormones have been implicated in modulating inflammation, with evidence suggesting that hormonal fluctuations can exacerbate inflammatory responses in females **(Monteiro et al. 2014; Collignon et al. 2024)**. In contrast, while male mice fed a WD showed more significant weight gain and adiposity, consistent with previous reports indicating a male-biased susceptibility to obesity-related metabolic changes, they did not show any diet-related visual impairments **(Samuelsson et al., 2008; Maric et al. 2022)**. Our findings revealed a divergence in neurovascular and metabolic phenotypes: males and females displayed increased retinal barrier breakdown and neuroinflammatory markers. However, males predominantly exhibited systemic metabolic disturbances, including increased visceral fat and elevated fasting glucose levels which females did not display. This dichotomy highlights the necessity of incorporating sex-based analyses into preclinical models to understand better the distinct pathways through which diet impacts health. These findings underscore the need to tailor therapeutic strategies to address sex-specific differences, focusing on neurovascular health in females and metabolic regulation in males.

Early-life WD exposure compromised BRB integrity and induced neuroinflammation, as evidenced by heightened microglial and astrocyte proliferation. This aligns with previous studies documenting similar inflammatory responses and vascular disruptions in adult mice exposed to prolonged WD **(Clarkson-Townsend et al., 2021; Boyé et al., 2022)**. In adult mice, 12 months of continued high-fat diet exposure is needed to observe similar effects in vascular density and BRB leakage **(Asare-Bediako et al. 2020; REF Rithwick Rajagopal 2015)**. Hence, our findings uniquely demonstrate that these changes manifest earlier (within just four months of WD exposure) when females are fed with this diet since first stages of development, emphasizing the heightened vulnerability of the developing retina to dietary insults during critical windows. Retinal microvasculature and BRB disruptions were apparent at this stage in female mice, even though functional tests such as the ERG revealed preserved functional responses. Other studies have shown small ERG changes specific to ondulatory potentials after 6 months of HFD feeding that correlated with glucose intolerance (**REF Rithwick Rajagopal** 2015). This suggests a latent phase where structural and inflammatory damage accumulates before functional impairments become detectable, likely requiring extended WD exposure to progress further. This disconnect highlights an urgent need for early interventions to mitigate damage before it evolves into irreversible functional decline. By mapping this trajectory from subclinical changes to overt dysfunction, future longitudinal studies could help identify critical time points for intervention, potentially preventing long-term complications like vision loss, associated with retinal dysfunction.

Moreover, our study revealed cognitive deficits and anxiety-like behaviors in WD-exposed female mice. Mice showed an impaired performance in the NOR test, a well-established method to assess recognition memory dependent on the hippocampus and perirhinal cortex **(Antunes et al. 2012)**. These findings align with previous studies linking maternal or early-life WD to cognitive dysfunctions in offspring, including deficits in memory, learning, and emotional regulation **(Cordner et al. 2019; Rodolaki et al. 2023)**. Our study adds evidence that visual processing deficits may contribute to cognitive impairments, offering a novel perspective on the interaction between sensory and cognitive systems. Emerging evidence suggests that retinal health can influence cognition due to shared neurovascular and inflammatory pathways **(Isceri et al. 2006; Trebbastoni et al. 2016; Casciano et al. 2024)**. For example, retinal dysfunction and reduced visual acuity in animal models correspond to deficits in spatial memory and anxiety regulation **(Brown et al. 2007; Storchi et al. 2019)**, further supporting a link between visual input and higher-order cognitive functions. In our study, WD-exposed female mice exhibited significant neuroinflammatory markers, paralleling findings that inflammation-induced disruptions in sensory systems can cascade into broader neurocognitive impairments. This interplay highlights a critical window during development when disruptions in sensory pathways, such as vision, can shape cognitive trajectories. By addressing these early sensory deficits through dietary interventions or targeted therapies, it may be possible to prevent the broader cognitive and emotional consequences observed in conditions like childhood obesity and metabolic syndrome.

Notably, obesity and metabolic disturbances during pregnancy have been implicated in developmental eye disorders **(Franzago et al., 2024)**. Maternal hyperglycemia, a hallmark of gestational diabetes and obesity, is known to impair early embryonic development, including ocular organogenesis, by inducing oxidative stress, inflammation, and epigenetic modifications in neural crest-derived tissues **(Lu et al., 2020; Wu et al., 2020).** Furthermore, hyperglycemia disrupts key signaling pathways such as Sonic Hedgehog (SHH) and Pax6, which are essential for optic vesicle development and lens differentiation **(Zhang et al., 2016; Cavodeassi et al., 2018)**. Our findings show a higher predisposition of females from maternal offspring to develop macroscopic abnormalities. These findings extend findings linking maternal hyperglycemia to disrupted optic vesicle and lens development **(Zhang et al. 2016, Lu et al. 2020)**. The observed female predisposition to these defects aligns with epidemiological trends in diet-related visual disorders, highlighting the need for sex-specific preventive strategies. A transcriptomic analysis at postnatal day 1 could reveal critical molecular signatures underlying these defects and further our understanding of their developmental origins.

Overall, this study underscores the critical importance of addressing sex-specific vulnerabilities in early-life dietary interventions. Advocating for proactive measures to mitigate dietary insults during critical developmental windows, preserving BRB integrity, and curbing early neuroinflammation may prevent retinal and cognitive health cascading effects. Moreover, our study provides a better model to understand the impact of dietary interventions on retinal health, reducing the exposure needed to these diets to observe mechanistic alterations associated with diabetic retinopathy.

Moving forward, several key areas warrant further investigation to fully understand the long-term impact of early-life WD exposure on retinal and cognitive health. <u>First,</u> longitudinal studies are essential to establish whether the observed structural and inflammatory changes progress into measurable functional deficits over time. <u>Second,</u> mechanistic insights into the molecular pathways underlying sex-specific effects of WD are crucial. Employing single-cell transcriptomics and epigenetic profiling **(Ying et al. 2021; Zibetti et al. 2022)** will help identify the key genes and signaling pathways involved in these differential responses. <u>Finally,</u> exploring therapeutic interventions is vital, particularly those aimed at restoring BRB integrity and attenuating neuroinflammation. Developing such therapies tailored to vulnerable populations, particularly females who appear more susceptible to WD-induced damage, could offer effective strategies for preventing or mitigating long-term cognitive and visual impairments.

Our findings emphasize the urgency of integrating dietary and lifestyle interventions into public health strategies, particularly during formative developmental periods when the neurovascular and cognitive systems are most vulnerable to environmental insults. The integration of such interventions could be vital to mitigating the growing burden of obesity-related visual and mental disorders in future generations.

## Materials and Methods

### Experimental Model

All experimental approaches were approved by the Yale University Animal Care and Use Committee protocol number 21043 and were by the National Institutes of Health guidelines. Adult mice (>8 weeks old) were used for all studies. Mice were housed in a 12-hour light–dark cycle (7:00–19:00) with ad libitum access to food and water unless otherwise indicated (fasting and WD studies). All experiments are in a wild-type (C57BL/6J) background (Jackson Laboratory 000664), Male and female mice were used for physiology studies. Western Diet used was from Research Diets (D12331) at 58% Fat and sucrose concentrations.

### Body weight and body composition

Body weights were monitored using a precision scale for 16 weeks. For feeding studies, mice were singly housed and acclimatized prior to the study. Daily food intake was manually measured using a precision scale. Blood samples were collected via the tail vein using a capillary collection system with EDTA (Sarstedt). Blood glucose concentration was measured using a glucometer (Aimstrip plus). Glucose tolerance tests (2 g/Kg) were performed on 4-hour fasted mice and blood glucose were measured at the indicated time-points. Whole-body composition was measured using NMR imaging (EchoMRI). Mice were restrained in a methacrylate restrainer and moved to the EchoMRI-500 body composition analysis device (EchoMRI). Three replicate measurements of fat mass and lean mass were taken. The average of the three replicates approximates the grams of fat and lean present in the animal. Measurements of core temperature were made using, an anal probe (Braintree Scientific).

### Open field test (OFT)

Open field test is a commonly used study for measuring locomotor activity and anxiety-like behavior. Our protocol was based on previous studies (Fan et al., 2019). Mice were place in the center of a dark methacrylate arena (40 x 40 cm) and allowed to freely explore it for 10 minutes. Trials were video recorded using Noldus camera equipment. Total distance, time spent in the center of the arena and time spent in the periphery was measured using the automated tracking software Ethovision XT version 17.5.

### Novel object recognition test (NORT)

The novel object recognition test is a highly employed cognitive tests for recognition memory. Our protocol was adapted from Leger et al., 2013 and Ramírez et al., 2022. The test was conducted in a methacrylate arena (40 x 40 cm) under an intensity light of 250 lux (measured with Light meter MT-912). The NORT consisted in three consecutive training days followed by a test phase. During the training, mice were exposed to two identical objects (Lego building blocks) once a day for a total time of 10 minutes for each session. These training sessions were performed for 3 consecutive days. After each session, mice were returned to the home cage. Arena and objects were cleaned with 70% ethanol to minimize olfactory cues between sessions.

At day 3, after the training session, mice were returned to the home cage for 1h before the text. In the test phase, mice were exposed to a new object (block of bigger size, different texture and color). Frequency and Cumulative time spent in the boundary zone of each object were measured using the tracking system Ethovision XT version 17.5. Additionally, discrimination indices were calculated as: (Time exploring novel object – Time exploring familiar object) / (Time exploring novel object + Time exploring familiar object).

### Elevated Plus Maze test (EPMT)

Mice were acclimatized to the room 1 week prior to the test and then placed individually on the central platform with their back to one of the open arms. Mice were tested for 5 minutes, during which they could freely explore the apparatus. Tracking software (Ethovision XT version 17.5) recognized mouse head, central body point, and the base of the tail. Anxiety was quantified by the frequency and accumulation time spent in the open arms. Higher anxiety is indicated by a lower frequency of movement into open arms and less time spend there.

### Barnes Maze test (BMT)

The Barnes maze test is a widely used cognitive test used to measure hippocampal-dependent spatial memory. Our protocol was based on previous studies (Ramírez et al., 2022). The maze consisted of a dark blue elevated circular platform (85cm stan height and 92 cm in diameter) with 20 equidistant holes located around the circumference. A black escape chamber used as a shelter for the mice, was placed underneath the designed target hole. The position of this target hole remained constant for each mouse during the acquisition phase. The reference cues were built with carton using different shapes and colors and were presented as walls surrounding the maze. Animals were tested under high-intensity light (superior to 20 Lux) to create an aversive environment on the surface of the apparatus, forcing them to explore the maze and find a refuge. The protocol consisted of 5 days of training, a two-day resting period, and the test day. On the first training day, a mouse was placed in the center of the maze and given 3 minutes to find the target hole and enter the attached escape chamber. In the case they did not enter on their own during the given time, the researcher gently placed the mouse to help them enter in the camber. Mice were allowed to stay there for 1 minute before returning them to the cage. This will enable the mouse to realize of the existence of the escape chamber and will give them practice to step down to the platform. The training was performed twice daily, with a 1-hour inter-trial interval, for 5 consecutive days. After the acquisition period, rodents typically remember the hole in which the escape chamber was placed and quickly proceed directly towards the hole. Improved performance over session reflects adequate learning. Mice were allowed to rest on days 6 and 7. The trial was performed on day 8. In this case, the escape chamber was removed and the animals were allowed to explore the maze for 2 minutes. During the test, the following parameters were scored: latency to target hole (defined as the time spent from the center of the circular platform to the precise hole where the escape chamber was located), time in quadrant (time spent in the quadrant of the platform where the escape hole was located) and the total distance (total distance traveled by the mouse during the 2-minute test). All trials were videos recorded. Analysis of the latency to find the target hole, time spent in the target quadrant and total distance were automatically measured by the video-tracking software.

### Brain staining

Animal brains were extracted intact and immersed in PFA overnight. Next, brains were sliced in the vibratome (Leica VT 1000S) to obtain 60 micras slices. Brains were stored in PBS at 4°C until use. A total of six brain slices were selected for each mouse including the areas of interest: Visual cortex (primary visual cortex V1) and Cingulate Cortex (cingulate cortex area A24a) from anterior to more posterior. Immunohistochemistry of cFOS in brain sections was performed as following: Brain slices were washed 4 times in PBS (10 mins per wash under soft agitation). The slices were blocked using a mix of 2% donkey serum in PBS and o.4% Triton X-100 for 1 hour. After that, brain slices were incubated in primary cFOS antibody (Cell Signaling Ref 06/2017) for 1 hour at room temperature and 48h in 4°C. They were washed 4 times in PBS (10 mins per wash under soft agitation). Following that, brain slices were incubated in secondary antibody (Alexa Fluor 568 Invitrogen A10042) at a concentration of 1:500. Brain slices were washed 4 times in PBS (10 mins per wash under soft agitation). Slices were mounted adding DAPI Fluoromont (SouthernBiotech Cat NO 010020) and covered with microscope cover glass for imaging.

### Retina dissection and immunostaining

Western diet and wildtype mice were deeply anesthetized with isoflurane and euthanized via cervical dislocation. Eyes were fixed in 4% formaldehyde at room temperature for 10 minutes. Retinas were isolated and stored in 100% methanol at −20°C overnight. The next day, the retinas were washed three times for 10 minutes each with PBS before incubation with specific primary antibodies. These antibodies were prepared in a blocking buffer containing 1% fetal bovine serum, 3% BSA, 0.5% Triton X-100, 0.01% sodium deoxycholate, and 0.02% sodium azide in PBS (pH 7.4). Incubation was performed overnight at 4°C with gentle shaking. On the following day, the retinas were washed again with three 10-minute PBS washes and incubated with secondary antibodies in buffer containing PBS pH 6.8, 1% Triton X-100, 0.1 mM CaCl_2_, 0.1 mM MgCl_2_, 0.1 mM MnCl_2_ at room temperature for 1 hour. Afterward, the retinas were washed three times with PBS, cut into four petals, and mounted using fluorescent mounting medium (DAKO, USA). The following antibodies were used: Claudin-5-GFP (Invitrogen, 352588), Plvap (BD Biosciences, 550563), Gfap (Abcam, ab4674), and Iba1 (Abcam, ab178846). IB4 and all secondary antibodies (donkey anti-primary, conjugated with Alexa Fluor 488, 568, or 647) were purchased from Invitrogen. Streptavidin-Texas Red (Vector Laboratories, SA-5006-1) was used to detect sulfo-NHS-biotin.

### Retro-orbitally tracer injection

Western diet and wildtype mice were anesthetized and then retro-orbitally injected with tracers and left to circulate for 30 minutes. The dose was modified according to the group: 1mg of sulfo-NHS-biotin/ mice in the case of the controls and 1.5 mg/ mice in the case of the obese animals (Thermo Scientific, 21217) was injected per mouse and 5mg of Cadaverine per gram of body weight (Thermo fisher A30679) was injected per mouse.

### Electroretinogram

Animals were dark adapted overnight in the same room where the test was taking place. Anesthesia was delivered before the measurement with a dose of 100mg/kg ketamine and 10mg/kg xylazine solution per hour. Anesthetized animals were placed on a stage with a heat pad to maintain appropriate body temperature during the test. The pupils were dilated with one drop per eye of 1% tropicamide solution. For scotopic (dim-light level) ERGs, single-flash responses were recorded at intensities of −4.0 to =2.7 log cd.s/m^2^. For photopic (bright-light level) ERGs, the mice were light-adapted under a background light of +1.48 log cd.s/m^2^ for 5-10 min. Single-flash photopic responses were recorded at intensities of −1.0 to +2.7 cd.s/m^2^, presented on the +1.48 log cd.s/m^2^ background. ERGs were recorded from both eyes simultaneously (Ganzfeld BigShot, LKC Technologies). ERG traces were analyzed using programs written in MATLAB (version R2024b, MathWorks). In brief, the ERG a-wave amplitude was calculated as the negative deflection observed withing the first 60 ms after the flash. The b-wave was calculated as the maximal amplitude of positive deflection following the a-wave after applying a 55 Hz Bessel filter to remove oscillatory potentials. Scotopic ERG b-wave amplitudes were fit using Equation 1, where Rmax,1 and Rmax,2 are the maximal response amplitudes and I0.5,1 and I0.5,2 are the half-saturating flash intensities; the first term reflects pure rod responses, whereas the second term reflects mixed rod/cone responses. We fit the data with a single term from Equation 1 for the scotopic a-wave. Data were calculated individually, and results are shown as mean ± SD:

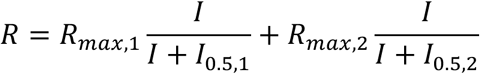

### Confocal microscopy and image analysis

Confocal images were captured using laser-scanning fluorescence microscopes (Zeiss LSM 900 and Leica SP8) with a 20X objective lens. Selective laser excitation at wavelengths of 405, 488, 547, or 647 nm was applied during imaging.

Claudin-5 analysis involved calculating the pixel intensity of thresholded vasculature relative to the corresponding pixel intensity in the thresholded IB4 channel. Sulfo-NHS-biotin and cadaverine leakage were measured as the mean pixel intensity, normalized to one control per sex group. Vascular density was quantified by measuring vascular area and mean on Ib4 staining for all groups and the normalized to one control per sex. Number of microglia were quantified by counting number of Iba1-positive cells per images, divided by vascular area. Astrocyte coverage was quantified by calculating the area of GFAP normalized by the area of IB4.

### Statistical Analysis

Data are expressed as mean ± SEM. Statistical analysis were performed using GraphPad Prism software. Two group-one factor comparisons were performed using a two-tailed unpaired Student’s t test. Datasets with two factors-one dependent variable were analyzed using two-way ANOVA followed by Sidak’s post-hoc test. One-sample t tests were performed to determine whether the NORT discrimination indexes observed in control/vehicle groups were significantly different from chance/0. In all cases p < 0.05. Symbols used are: ^∗^p < 0.05; ^∗∗^p < 0.01; ^∗∗∗^p < 0.001; ^∗∗∗∗^p < 0.0001. Statistical parameters can be found in the Figures and Figure legends. For analysis of ERG data, we focused on intensity ranges that approximate the maximal rod-only response and mixed rod/cone response based on the apparent saturation of each response level according to the fit with Equation 1. Scotopic a-waves and photopic b-waves were analyzed only at the higher intensity range (+2 to +2.7 log cd·s/m2), where the signal-to-noise ratio was sufficient. In Figure 3, where three genotypes were compared to control, we reduced the α in unpaired t tests by ten-fold to account for multiple comparisons (from 0.05 to 0.005). For analysis of GFAP data, images were processed with Fiji (Version: 2.14.0-1.54f) to calculate the area and the mean of GFAP positive signal. This was normalized by dividing the value of GFAP by the value of IB4 for each sample. An unpaired t-test was performed to compared control group against western diet group. A value lower than 0.05 was considered statistically significant. For analysis of cFOS, positive cells were manually counted by trained staff. Each brain slice was quantified twice, including each hemisphere, and they were all added together to have a representative value of each brain loci.

## Acknowledgements

M.S. acknowledges support from the McCluskey family and Interstellar Initiative (NYAS/AMED). This work was supported by the National Institute of Diabetes, Digestive and Kidney Diseases R00DK1208689, DRC P30 DK045735 and National Institute of Aging P30AGO66508. J-LT declares research grant support from the US National Institutes of Health and Monitoring Neuroinflammation supported by the Fonds Recherche Neurosciences.

## Author contributions

M.S. and D.M. conceived and designed the study and developed the research program. D.M., J.F., T.Z., A.R., J.G.R., and D.N. performed experiments. J.D. conducted the ERG study design and helped interpret the results. A.E. conducted the retina immunofluorescent study designs. M.S. wrote the manuscript with input from all the authors.

## References

Blüher M. Obesity: global epidemiology and pathogenesis. Nat Rev Endocrinol. 2019 May;15(5):288-298. doi: 10.1038/s41574-019-0176-8. PMID: 30814686 Saltiel AR, Olefsky JM. Inflammatory mechanisms linking obesity and metabolic disease. J Clin Invest. 2017 Jan 3;127(1):1–4. doi: 10.1172/JCI92035. Epub 2017 Jan 3. PMID: 28045402; PMCID: PMC5199709.

Antonetti DA, Klein R, Gardner TW. Diabetic retinopathy. N Engl J Med. 2012 Mar 29;366(13):1227-39. doi: 10.1056/NEJMra1005073. PMID: 22455417.

Catalano PM, Shankar K. Obesity and pregnancy: mechanisms of short term and long term adverse consequences for mother and child. BMJ. 2017 Feb 8;356:j1. doi: 10.1136/bmj.j1. PMID: 28179267; PMCID: PMC6888512.

WHO, 2023: https://www.who.int/news-room/fact-sheets/detail/obesity-and-overweight

CDC, 2023: https://www.cdc.gov/obesity/childhood-obesity-facts/childhood-obesity-facts.html

Rundle AG, Park Y, Herbstman JB, Kinsey EW, Wang YC. COVID-19-Related School Closings and Risk of Weight Gain Among Children. Obesity (Silver Spring). 2020 Jun;28(6):1008–1009. doi: 10.1002/oby.22813. Epub 2020 Apr 18. PMID: 32227671; PMCID: PMC7440663 Dezor-Garus J, Niechciał E, Kędzia A, Gotz-Więckowska A. Obesity-induced ocular changes in children and adolescents: A review. Front Pediatr. 2023 Mar 23;11:1133965. doi: 10.3389/fped.2023.1133965. PMID: 37033164; PMCID: PMC10076676.

Pacifico, L., Perla, F. M., & Chiesa, C. (2020). Nonalcoholic fatty liver disease in children: current and novel insights into biomarkers, pathogenesis, and therapeutic approaches. Pediatric Research, 87(3), 456–463. doi:10.1038/s41390-019-0581-5

Travers SH, Labarta JI, Gargosky SE, Rosenfeld RG, Jeffers BW, Eckel RH. Insulin-like growth factor binding protein-I levels are strongly associated with insulin sensitivity and obesity in early pubertal children. J Clin Endocrinol Metab. 1998 Jun;83(6):1935–9. doi: 10.1210/jcem.83.6.4857. PMID: 9626122.

Stanek A, Brożyna-Tkaczyk K, Myśliński W. The Role of Obesity-Induced Perivascular Adipose Tissue (PVAT) Dysfunction in Vascular Homeostasis. Nutrients. 2021 Oct 28;13(11):3843. doi: 10.3390/nu13113843. PMID: 34836100; PMCID: PMC8621306.

Keeling E, Lynn SA, Koh YM, Scott JA, Kendall A, Gatherer M, Page A, Cagampang FR, Lotery AJ, Ratnayaka JA. A High Fat “Western-style” Diet Induces AMD-Like Features in Wildtype Mice. Mol Nutr Food Res. 2022 Jun;66(11):e2100823. doi: 10.1002/mnfr.202100823. Epub 2022 Apr 28. PMID: 35306732; PMCID: PMC9287010.

Selvam S, Kumar T, Fruttiger M. Retinal vasculature development in health and disease. Prog Retin Eye Res. 2018 Mar;63:1–19. doi: 10.1016/j.preteyeres.2017.11.001. Epub 2017 Nov 10. PMID: 29129724.

Vogt MC, Paeger L, Hess S, Steculorum SM, Awazawa M, Hampel B, Neupert S, Nicholls HT, Mauer J, Hausen AC, Predel R, Kloppenburg P, Horvath TL, Brüning JC. Neonatal insulin action impairs hypothalamic neurocircuit formation in response to maternal high-fat feeding. Cell. 2014 Jan 30;156(3):495–509. doi: 10.1016/j.cell.2014.01.008. Epub 2014 Jan 23. PMID: 24462248; PMCID: PMC4101521.

Samuelsson AM, Matthews PA, Argenton M, Christie MR, McConnell JM, Jansen EH, Piersma AH, Ozanne SE, Twinn DF, Remacle C, Rowlerson A, Poston L, Taylor PD. Diet-induced obesity in female mice leads to offspring hyperphagia, adiposity, hypertension, and insulin resistance: a novel murine model of developmental programming. Hypertension. 2008 Feb;51(2):383–92. doi: 10.1161/HYPERTENSIONAHA.107.101477. Epub 2007 Dec 17. PMID: 18086952.

Franzago M, Borrelli P, Di Nicola M, Cavallo P, D’Adamo E, Di Tizio L, Gazzolo D, Stuppia L, Vitacolonna E. From Mother to Child: Epigenetic Signatures of Hyperglycemia and Obesity during Pregnancy. Nutrients. 2024 Oct 16;16(20):3502. doi: 10.3390/nu16203502. PMID: 39458497; PMCID: PMC11510513.

Wenhui Lu; Peixin Yang. June 2020 1334-P: Hyperglycemia-Induced Eye Malformations through Dysregulation of Autophagy in the Mouse Embryo. 10.2337/db20-1334-P

Wu Y, Liu B, Sun Y, Du Y, Santillan MK, Santillan DA, Snetselaar LG, Bao W. Association of Maternal Prepregnancy Diabetes and Gestational Diabetes Mellitus With Congenital Anomalies of the Newborn. Diabetes Care. 2020 Dec;43(12):2983–2990. doi: 10.2337/dc20-0261. Epub 2020 Oct 21. PMID: 33087319; PMCID: PMC7770264.

Zhang SJ, Li YF, Tan RR, Tsoi B, Huang WS, Huang YH, Tang XL, Hu D, Yao N, Yang X, Kurihara H, Wang Q, He RR. A new gestational diabetes mellitus model: hyperglycemia-induced eye malformation via inhibition of Pax6 in the chick embryo. Dis Model Mech. 2016 Feb;9(2):177–86. doi: 10.1242/dmm.022012. Epub 2016 Jan 7. PMID: 26744353; PMCID: PMC4770145.

Cavodeassi F, Creuzet S, Etchevers HC. The hedgehog pathway and ocular developmental anomalies. Hum Genet. 2019 Sep;138(8-9):917–936. doi: 10.1007/s00439-018-1918-8. Epub 2018 Aug 2. PMID: 30073412; PMCID: PMC6710239.

American Academy of Ophtalmology 2024 Release: https://www.aao.org/newsroom/news-releases/detail/women-face-higher-risk-of-blindness-than-men

Rice D, Barone S Jr. Critical periods of vulnerability for the developing nervous system: evidence from humans and animal models. Environ Health Perspect. 2000 Jun;108 Suppl 3(Suppl 3):511-33. doi: 10.1289/ehp.00108s3511. PMID: 10852851; PMCID: PMC1637807.

Rudraraju M, Narayanan SP, Somanath PR. Regulation of blood-retinal barrier cell-junctions in diabetic retinopathy. Pharmacol Res. 2020 Nov;161:105115. doi: 10.1016/j.phrs.2020.105115. Epub 2020 Aug 1. PMID: 32750417; PMCID: PMC7755666.

Eltanani S, Yumnamcha T, Gregory A, Elshal M, Shawky M, Ibrahim AS. Relative Importance of Different Elements of Mitochondrial Oxidative Phosphorylation in Maintaining the Barrier Integrity of Retinal Endothelial Cells: Implications for Vascular-Associated Retinal Diseases. Cells. 2022 Dec 19;11(24):4128. doi: 10.3390/cells11244128. PMID: 36552890; PMCID: PMC9776835.

Boyé K, Geraldo LH, Furtado J, Pibouin-Fragner L, Poulet M, Kim D, Nelson B, Xu Y, Jacob L, Maissa N, Agalliu D, Claesson-Welsh L, Ackerman SL, Eichmann A. Endothelial Unc5B controls blood-brain barrier integrity. Nat Commun. 2022 Mar 4;13(1):1169. doi: 10.1038/s41467-022-28785-9. PMID: 35246514; PMCID: PMC8897508.

Yao X, Zhao Z, Zhang W, Liu R, Ni T, Cui B, Lei Y, Du J, Ai D, Jiang H, Lv H, Li X. Specialized Retinal Endothelial Cells Modulate Blood-Retina Barrier in Diabetic Retinopathy. Diabetes. 2024 Feb 1;73(2):225–236. doi: 10.2337/db23-0368. PMID: 37976214.

Kim SY, Cheon J. Senescence-associated microvascular endothelial dysfunction: A focus on the blood-brain and blood-retinal barriers. Ageing Res Rev. 2024 Sep;100:102446. doi: 10.1016/j.arr.2024.102446. Epub 2024 Aug 5. PMID: 39111407.

Li Y, Wang C, Zhang L, Chen B, Mo Y, Zhang J. Claudin-5a is essential for the functional formation of both zebrafish blood-brain barrier and blood-cerebrospinal fluid barrier. Fluids Barriers CNS. 2022 Jun 3;19(1):40. doi: 10.1186/s12987-022-00337-9. PMID: 35658877; PMCID: PMC9164509.

Noailles A, Fernández-Sánchez L, Lax P, Cuenca N. Microglia activation in a model of retinal degeneration and TUDCA neuroprotective effects. J Neuroinflammation. 2014 Oct 29;11:186. doi: 10.1186/s12974-014-0186-3. PMID: 25359524; PMCID: PMC4221719.

Hanke-Gogokhia C, Zapadka TE, Finkelstein S, Klingeborn M, Maugel TK, Singer JH, Arshavsky VY, Demb JB. The Structural and Functional Integrity of Rod Photoreceptor Ribbon Synapses Depends on Redundant Actions of Dynamins 1 and 3. J Neurosci. 2024 Jun 19;44(25):e1379232024. doi: 10.1523/JNEUROSCI.1379-23.2024. PMID: 38641407; PMCID: PMC11209669.

Antunes M, Biala G. The novel object recognition memory: neurobiology, test procedure, and its modifications. Cogn Process. 2012 May;13(2):93–110. doi: 10.1007/s10339-011-0430-z. Epub 2011 Dec 9. PMID: 22160349; PMCID: PMC3332351.

Rodríguez Peris L, Scheuber MI, Shan H, Braun M, Schwab ME. Barnes maze test for spatial memory: A new, sensitive scoring system for mouse search strategies. Behav Brain Res. 2024 Feb 26;458:114730. doi: 10.1016/j.bbr.2023.114730. Epub 2023 Oct 26. PMID: 37898351.

Keleher MR, Zaidi R, Patel K, Ahmed A, Bettler C, Pavlatos C, Shah S, Cheverud JM. The effect of dietary fat on behavior in mice. J Diabetes Metab Disord. 2018 Nov 22;17(2):297–307. doi: 10.1007/s40200-018-0373-3. PMID: 30918865; PMCID: PMC6405378.

Tsan L, Décarie-Spain L, Noble EE, Kanoski SE. Western Diet Consumption During Development: Setting the Stage for Neurocognitive Dysfunction. Front Neurosci. 2021 Feb 10;15:632312. doi: 10.3389/fnins.2021.632312. PMID: 33642988; PMCID: PMC7902933.

Hayes AMR, Lauer LT, Kao AE, Sun S, Klug ME, Tsan L, Rea JJ, Subramanian KS, Gu C, Tanios N, Ahuja A, Donohue KN, Décarie-Spain L, Fodor AA, Kanoski SE. Western diet consumption impairs memory function via dysregulated hippocampus acetylcholine signaling. Brain Behav Immun. 2024 May;118:408–422. doi: 10.1016/j.bbi.2024.03.015. Epub 2024 Mar 8. PMID: 38461956; PMCID: PMC11033683.

López-Taboada I, González-Pardo H, Conejo NM. Western Diet: Implications for Brain Function and Behavior. Front Psychol. 2020 Nov 23;11:564413. doi: 10.3389/fpsyg.2020.564413. PMID: 33329193; PMCID: PMC7719696.

Clarkson-Townsend DA, Douglass AJ, Singh A, Allen RS, Uwaifo IN, Pardue MT. Impacts of high fat diet on ocular outcomes in rodent models of visual disease. Exp Eye Res. 2021 Mar;204:108440. doi: 10.1016/j.exer.2021.108440. Epub 2021 Jan 11. PMID: 33444582; PMCID: PMC7946735.

